# Verbenone and L-arginine from *Daucus carota* Seeds Attenuate Oxidative Stress in Streptozotocin-Nicotinamide (STZ-NAD)-Induced Diabetic Mice: Evidence from *In vitro* and *In vivo* Studies

**DOI:** 10.1101/2023.09.28.559552

**Authors:** Habibu Tijjani, Umar Ibrahim, Sadiq Tijani, Sadiya Alka, Ahmed Olatunde, Aminu Umar Kura, Haladu Ali Gagman, Oluremi A. Saliu, Oluwafemi A. Idowu, Frank Eromosele Ebhodaghe

## Abstract

Free radicals are produced in diabetes mellitus (DM), due to protein glycosylation and glucose autooxidation. However, compounds from plants were proven to be effective in the management of DM and related conditions. This study evaluated the antioxidant effect of verbenone, L-arginine, and their ratio combinations in diabetic mice. DM was induced in mice by intraperitoneal (IP) injection of streptozotocin and nicotinamide adenine dinucleotide (NAD) and the diabetic mice were treated with verbenone, L-arginine, and their ratios for 28 days. In the *in vitro* study, L-arginine expressed higher 1,1-diphenyl-2-picrylhydrazyl (DPPH) radical scavenging activity, while verbenone and L-arginine expressed higher nitric oxide (NO) and hydrogen peroxide (H_2_O_2_) scavenging activities, especially at higher concentrations when compared with vitamin C. At the end of the treatment period, the levels of blood glucose, total protein, and malondialdehyde were significantly increased while the levels of reduced glutathione, nitrite, and activities of glutathione peroxidase, glutathione transferase, catalase, superoxide dismutase were significantly decreased in the diabetic untreated mice. However, these diabetes-induced alterations were reversed to normal levels after the administration of verbenone, L-arginine, and their ratio combinations at 100 and 200 mg/kg body weight. Furthermore, *in silico* studies revealed the antioxidant potential of both verbenone and L-arginine by their interaction with antioxidant proteins, expressing their potential antioxidant properties. The results of the study indicated that verbenone, L-arginine, and their ratio combination possess antioxidant property and attenuate oxidative stress in diabetic mice.

**Highlights:**

- Verbenone and L-arginine are natural compounds found in *Daucus carota* seeds and other plants.
- Verbenone and L-arginine possess *in vitro* and *in vivo* antioxidant activities.
- Verbenone, L-arginine and their ratio combination (1:1) enhance the activities of antioxidant enzymes in streptozotocin-nicotinamide (NAD-STZ) induced diabetic mice.
- Furthermore, the two compounds interacted with antioxidant proteins, expressing their potential antioxidant property in an *in silico* model.

## 1. Introduction

Antioxidants are those substances that inhibit and stabilized radicals from damaging tissues through donation of electrons to these reactive radicals. Intake of antioxidant-rich foods and vegetables is reported to reduce the risk of numerous diseases associated with oxidative stress (Hamid *et al*., 2010). Several disorders such as cancer, diabetes mellitus (DM), cardiovascular and neurodegenerative diseases, are associated with the accumulation of unstable radicals, which inturn leads to oxidative stress (Gilgun-sherk *et al*., 2002; Islam *et al*., 2013). Attention has been drawn to the intake of foods and vegetables rich in natural antioxidants, to confer protection on the body against the damaging effects of free reactive radicals. Thus, they could be vital as nutraceuticals and functional foods against several oxidative stress-related diseases

*Daucus carota* are well-known vegetables usually consumed globally and with many health promoting properties. Some studies revealed that the seeds from this plant are medicinally important in the management of DM through the inhibition of oxidative stress (Pouraboli and Ranjbar, 2015; Tijjani *et al*., 2019; Tijjani *et al*., 2020a; 2020b; 2020c; Tijjani and Imam, 2021). Verbenone and L-arginine are examples of compounds found in *D. carota* seed. L-arginine (2- amino-5-guanodino-pentenoic acid, Figure 1a) is an important amino acid in human and animal health and growth. The deficiency of L-arginine in infants causes hyperammonemia and multiple organ abnormalities (Wu *et al*., 2004). Also, a reduction in plasma concentration of arginine was reported to be associated with DM (Pieper *et al*., 1996). To further buttress its role in DM, L-arginine plays an important role in the metabolism of nitric oxide (NO), through its rich nitrogen atom, and therefore plays a metabolic regulatory role via nitric oxide (Jobgen *et al*., 2006). Therefore, the amino acid regulates insulin secretion through NO-dependent and NO-independent pathways (Schmidt *et al*., 1992; Thams and Capito, 1999).

**Figure 1:**
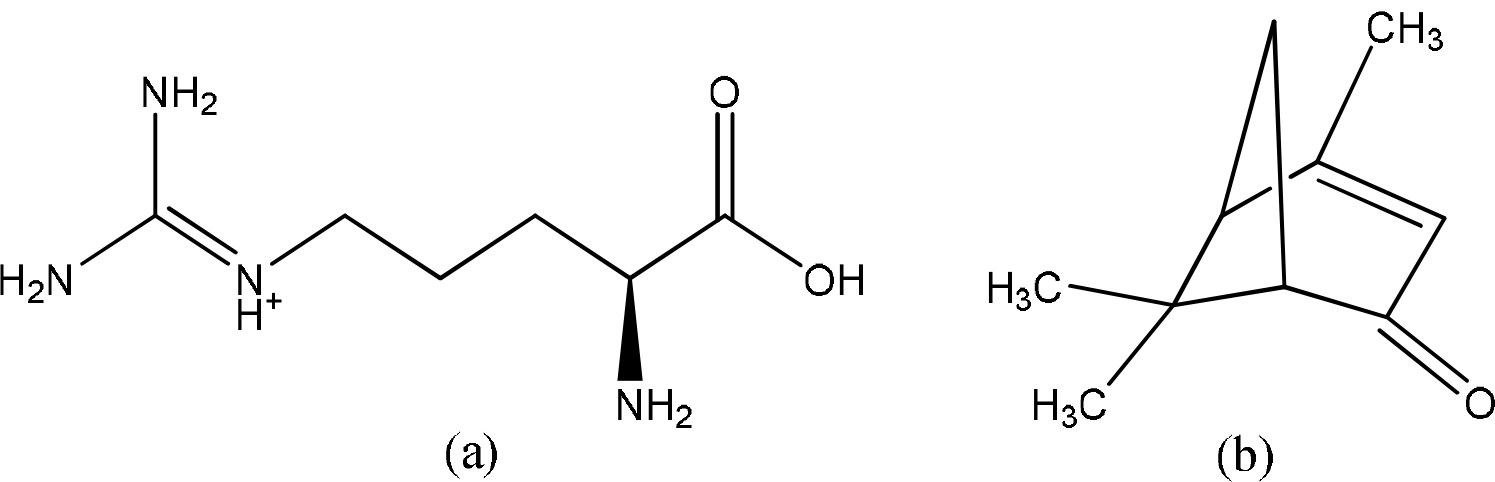
Chemical structures of (a) L-arginine and (b) verbenone

Verbenone (trimethyl-bicyclo-heptenone, Figure 1b) is a natural bicyclic ketone mono-terpene with several pharmacological properties including anti-aggregation pheromone (Kostyk *et al*., 1993), repellent (Miller *et al*., 1995; Zhang *et al*., 2006), anti-inflammatory, bronchodilator and haemolytic actions (Rappaport *et al*., 2001). Verbenone oxime ester, an example of verbenone derivative, was discovered to display anticancer (Song *et al*., 2005), antifungal (Wang *et al*., 2014), antioxidant (Sammaiah *et al*., 2015; Harini *et al*., 2012), antiviral (Ouyang *et al*., 2008) properties. The search for natural antioxidants to attenuate oxidative stress-related diseases such as DM is on the increase and plants such as vegetables are gaining more attention. Therefore, the present study was designed to evaluate the *in vitro* and *in vivo* antioxidant activities of L-arginine and verbenone in streptozotocin-nicotinamide adenine dinucleotide (STZ-NAD)-induced diabetic mice as well as the effects of their ratio combination (1:1 v/m), since their combination could improve their solubilities (Tijjani, 2022). Furthermore, the binding interactions of the compounds in antioxidant proteins were evaluated using an *in silico* model.

## 2. Material and methods

### 2.1. Chemicals

Verbenone, L-arginine, 1,1-diphenyl-2-picrylhydrazyl (DPPH), streptozotocin (STZ), nicotinamide adenine dinucleotide (NAD), and dimethyl sulfoxide (DMSO) were purchased from Carbosynth Limited, UK. Methanol, sodium phosphate, ammonium molybdate, sodium nitroprusside, hydrogen peroxide, and FeCl_3_ were purchased from Sigma Aldrich, Germany. All other chemicals were of analytical grades.

### 2.2. *In vitro* antioxidant analysis

*In vitro* antioxidant analyses was performed according to the methods of McCune and Johns, (2002), Prieto *et al*. (1999), Fiorentino *et al*. (2008), Ruch *et al*. (1989), and Girgih *et al*. (2013) for 1,1-diphenyl-2-picrylhydrazyl (DPPH) scavenging activity, total antioxidant capacity (TAC), nitric oxide (NO) scavenging activity, hydrogen peroxide (H_2_O_2_) scavenging activity and ferric reducing antioxidant power (FRAP) assays of the compounds (Verbenone, L-arginine, and their ratios) respectively.

### 2.3. Experimental animals

Wistar albino mice were obtained from the animal house of the Department of Pharmacology, Bauchi State University Gadau, Nigeria. The mice were kept under laboratory conditions with 12h day and night cycles. The experimental mice were fed on animal feed, and water *ad libitum*. The research protocol was approved by the Ethical Review Committee of the University of Jos, Nigeria with the ethical clearance number UJ/FPS/F17-00379.

### 2.4. Induction of diabetes

Diabetes was induced in overnight fasted mice by a single intraperitoneal injection of STZ at a dose of 60 mg/kg body weight in 0.1 M citrate buffer (pH 4.5), 10 minutes after injection with 110 mg/kg body weight NAD solution prepared in normal saline (pH 4.5). The mice were given 10% dextrose after 6hr of STZ administration to prevent STZ-induced hypoglycemia. The induction of diabetes was verified after 72 hr by measuring blood glucose levels using glucose test strips (Accu-Check) and mice with blood glucose values of 200 mg/dL and above were considered diabetic and were used for the study.

### 2.5. Animal grouping and administration

The animals were randomly distributed into nine (8) groups containing five (5) mice each. Group I (Control) are none diabetic + none treated mice, which received 5% DMSO, group II (Diabetic) are diabetic untreated mice, which received 5% DMSO, group III (Glibenclamide) are diabetic mice treated with 5 mg/kg bwt glibenclamide, group IV (Verbenone 100) are diabetic mice treated with 100 mg/kg bwt verbenone, group V (L-arginine 100) are diabetic mice treated with 100 mg/kg bwt L-arginine, group VI (L-arginine 200) are diabetic mice treated with 200 mg/kg bwt L-arginine, group VII (Ver-L-Arg 100) are diabetic mice treated with 100 mg/kg bwt of verbenone + L-arginine ratio (1:1 v/m), while group VIII (Ver-L-Arg 200) are diabetic mice treated with 200 mg/kg bwt of verbenone + L-arginine (1:1 v/m). All doses were prepared in 5% DMSO and administered in 0.2 mL per dosage. Treatments were done once daily for 14 days, after which the mice were sacrificed under anesthesia (diethyl ether). Blood and liver tissue samples were collected for subsequent analyses.

### 2.6. Collection and preparation of blood and liver homogenates

All experimental animals were weighed and sacrificed on the 15^th^ day, 24 hr after the last intervention period, and blood samples from the mice were collected into EDTA containers. Plasma was separated from the collected blood samples by centrifugation at 1500 x g for 10 min into appropriately labeled test tubes for various analyses. Liver was excised from the experimental mice, cleaned, and weighed. The liver was then homogenized using 0.25 M sucrose solution (1:5 w/v), then centrifuged at 1500 x g for 5 min to obtain supernatants that were appropriately collected and labeled in test tubes for subsequent analyses.

### 2.7. Determination of biochemical parameters

The total protein was measured according to the method of Bradford (1976). Reduced glutathione (GSH), nitric oxide (NO), and malondialdehyde (MDA) concentrations were determined according to the methods of Beutler *et al*. (1963), Green *et al*. (1982), and Varshney and Kale (1990) respectively. The methods described by Habig *et al*. (1974), Misra and Fridovich (1972), Rotruck *et al*. (1973), and Sinha (1972) were used to determine glutathione-S-transferase (GST), superoxide dismutase (SOD), glutathione peroxidase (GPx) and catalase (CAT) respectively.

### 2.8. Molecular docking studies

Chemical structures of verbenone (PubChem CID 29025), L-arginine (PubChem CID 6322), 5- fluorouracil (FLU, PubChem CID 3385), zileuton (ZIL, PubChem CID 60490), melatonin (MEL, PubChem CID 896), dextromethorphan (DEX, PubChem CID 5360696) were retrieved from PubChem database (www.pubchem.ncbi.nlm.nih.gov) (Kim *et al*., 2019). They were converted to pdbqt format after setting an appropriate torsion center for each ligand.

Crystal structures of human cytochrome P450 (CP450, 10G5) (Williams *et al*., 2003), lipoxygenase in complex with protocatechuic acid (LO, 1N8Q) (Borbulevych *et al*., 2004), human myeloperoxidase-thiocyanate complex (MP, 1DNU) (Blair-Johnson *et al*., 2001) and NAD(P)H oxidase from *Lactobacillus sanfranciscensis* (NAD(P)HO, 2CDU) (Lountos *et al*., 2006) were downloaded from the protein databank (www.rcsb.org) (Berman *et al*., 2000) with their respective PDB codes. They were prepared by removing existing ligands and water molecules, while missing polar hydrogen atoms were added using Autodock v4.2 program, Scripps Research Institute. They were saved in dockable pdbqt format for molecular docking studies.

The molecular docking studies of verbenone, L-arginine, the reference ligands, and protein targets were carried out using Autodock Vina (Trott and Olson, 2010). The binding energies of each compound were recorded while the interactions were analyzed and viewed using Discovery Studio Visualizer, BIOVIA, 2020.

### 2.9. Statistical analysis

Data were expressed as mean ± Standard Error of Mean (SEM) where applicable. They were subjected to Analysis of Variance (ANOVA) followed by Dunnett’s test using SPSS version 20, PSS Inc., Chicago. IL, USA. Significance was considered at p<0.05. Graphs were plotted using GraphPad Prism 6 software (GraphPad Software, California, USA).

## 3. Results

### 3.1. *In vitro* antioxidants activities verbenone, L-arginine and their ratio (Ver-L-Arg)

The DPPH radical scavenging activities of verbenone, L-arginine, and their ratio combination (1:1) were comparable with that of vitamin C (Figure 2a). The DPPH radical scavenging activities of verbenone, and the ratio combination (Ver-L-Arg) were significantly lower than that of L-arginine. However, verbenone, L-arginine and their ratio combination (Ver-L-Arg) expressed significantly higher NO scavenging activities compared with vitamin C (Figure 2b). The TAC of verbenone, L-arginine and their ratio combination (Ver-L-Arg) were significantly lower compared with vitamin C (Figure 2c), except at higher concentration (100 μg/mL), where the activity was not significantly different in verbenone and L-arginine. Similar results were observed in H_2_O_2_ scavenging activities and ferric ion reducing properties of verbenone, L-arginine, and the ratio combination (Ver-L-Arg) (Figure 2d – 2e). At higher concentrations, verbenone expressed higher scavenging activity and reducing property compared with vitamin C.

**Figure 2.**
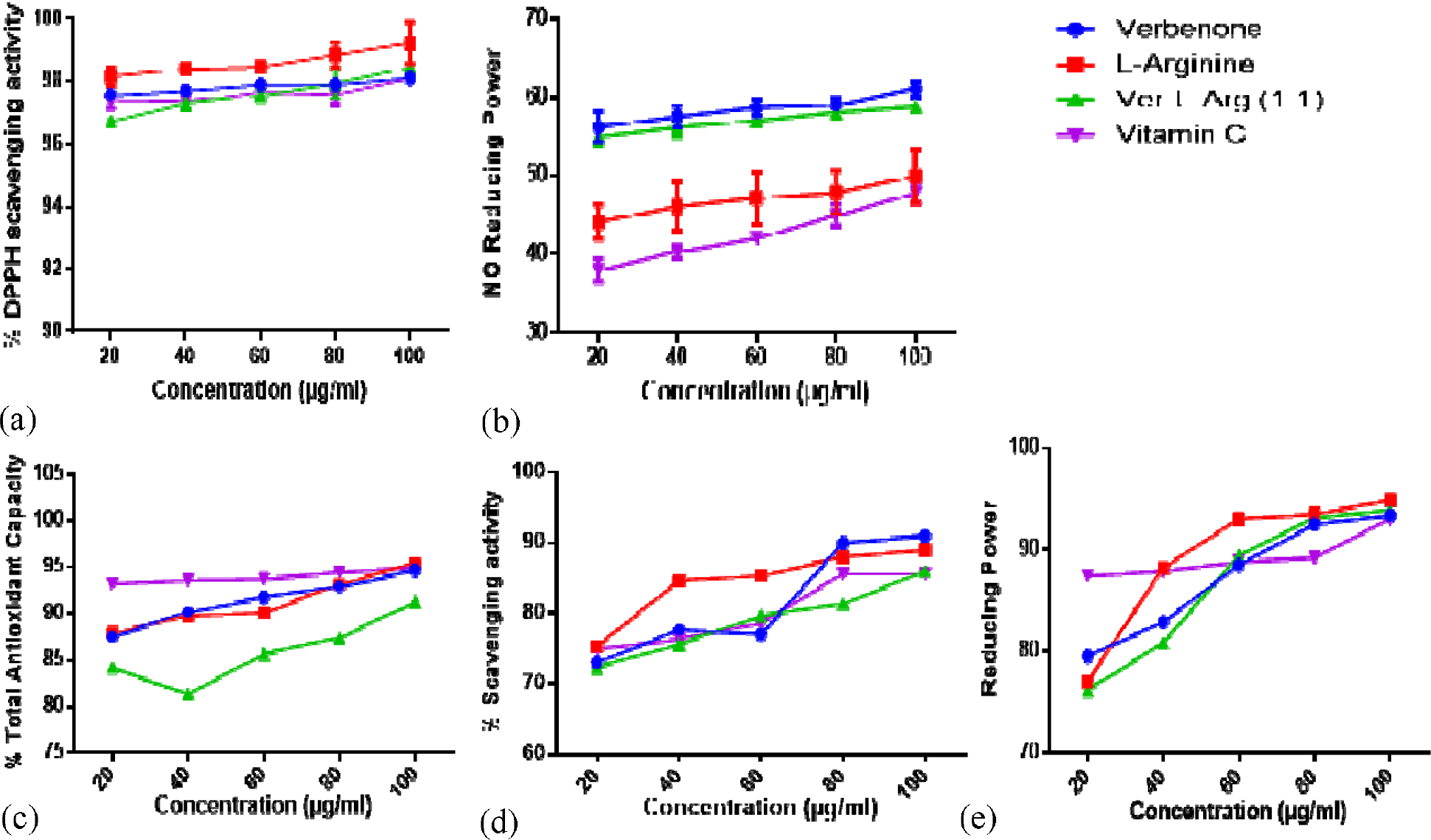
*In vitro* antioxidant activites of verbenone, L-arginine and their ratio combination. (a) DPPH scavenging activity (b) NO scavenging capacity (c) TAC (d) % H_2_O_2_ scavenging activity (e) FRAP. Values were expressed as means ± SEM of triplicate determination. Ver-L-Arg (1:1) = verbenone + L-arginine ratio (1:1 v/m)

### 3.2. Effects of verbenone and L-arginine in STZ-NAD-induced diabetic mice

The diabetic untreated mice showed significant (p<0.05) increase in blood glucose level (533.00) compared to the control mice (120.00) (Table 1). However, all the diabetic treated mice showed decrease in blood glucose levels compared to the diabetic untreated mice, except L-arginine treatment at 100 mg/kg bwt. Also, treatment with glibenclamide (5 mg/kg bwt) showed reduced blood glucose level from 216.00 to 158.00 mg/dL. Furthermore, Ver-L-Arg 200 group (38.42%) and verbenone 100 group (38.23%) displayed the highest blood glucose glucose lowering effects compared with glibenclamide (5 mg/kg bwt) treated group with 45.70% glycemic reduction.

**Table 1:** Effect of verbenone, L-arginine and their ratio combinaiton on blood glucose level in STZ-NAD-induced diabetic mice.

| Treatment groups | Blood glucose level (mg/dL) |  |  |  |
| --- | --- | --- | --- | --- |
|  | Day 0 | Day 4 | Day 10 | Day 15 |
| Control | 127.00±8.00 | 115.00±8.00 | 137.00±4.00 | 120.00±6.00 |
| Diabetic Control | 315.00±6.00 | 479.00±8.00 | 586.00±19.00 | 533.00±11.00 |
| Glibenclamide 5 mg/kg bwt | 316.00±6.00 | 265.00±11.00 | 224.00±6.00 | 158.00±23.00 |
| Verbenone 100 mg/kg bwt | 395.00±7.00 | 165.00±7.00 | 250.00±12.00 | 244.00±11.00 |
| L-arginine 100 mg/kg bwt | 356.00±3.00 | 286.00±7.00 | 512.00±16.00 | 521.00±25.00 |
| L-arginine 200 mg/kg bwt | 346.00±4.00 | 167.00±3.00 | 535.00±18.00 | 424.00±10.00 |
| Ratio (1:1 v/m) 100 mg/kg bwt | 413.00±5.00 | 318.00±9.00 | 560.00±23.00 | 511.00±11.00 |
| Ratio (1:1 v/m) 200 mg/kg bwt | 367.00±4.00 | 147.00±5.00 | 207.00±10.00 | 226.00±5.00 |

Values are expressed as mean ± SEM (n = 5). Ratio (1:1 v/m) = Verbenone + L-arginine (1:1 v/m)

### 3.3. Effects of verbenone, L-arginine and their ratio combination on non-enzymatic antioxidant indices

The plasma total protein was significant increase (p<0.05) in the diabetic untreated and Ver-L-Arg 200 groups compared with control group (Figure 3a), with a significant decrease (p<0.05) in L-arginine treatments at 100 and 200 mg/kg body weights. However, in the liver, a significant decrease (p<0.05) was recorded in glibenclamide, L-arginine 100, and Ver-L-Arg 200 groups compared to control group (Figure 3b). The levels of reduced glutathione in plasma (pGSH) was significantly reduced (p<0.05) in diabetic untreated and Ver-L-Arg 200 groups compared with control group. But, no significant difference (p>0.05) was observed in liver GSH (lGSH) in all treated groups except in Ver-L-Arg 200 group compared to control mice.

**Figure 3.**
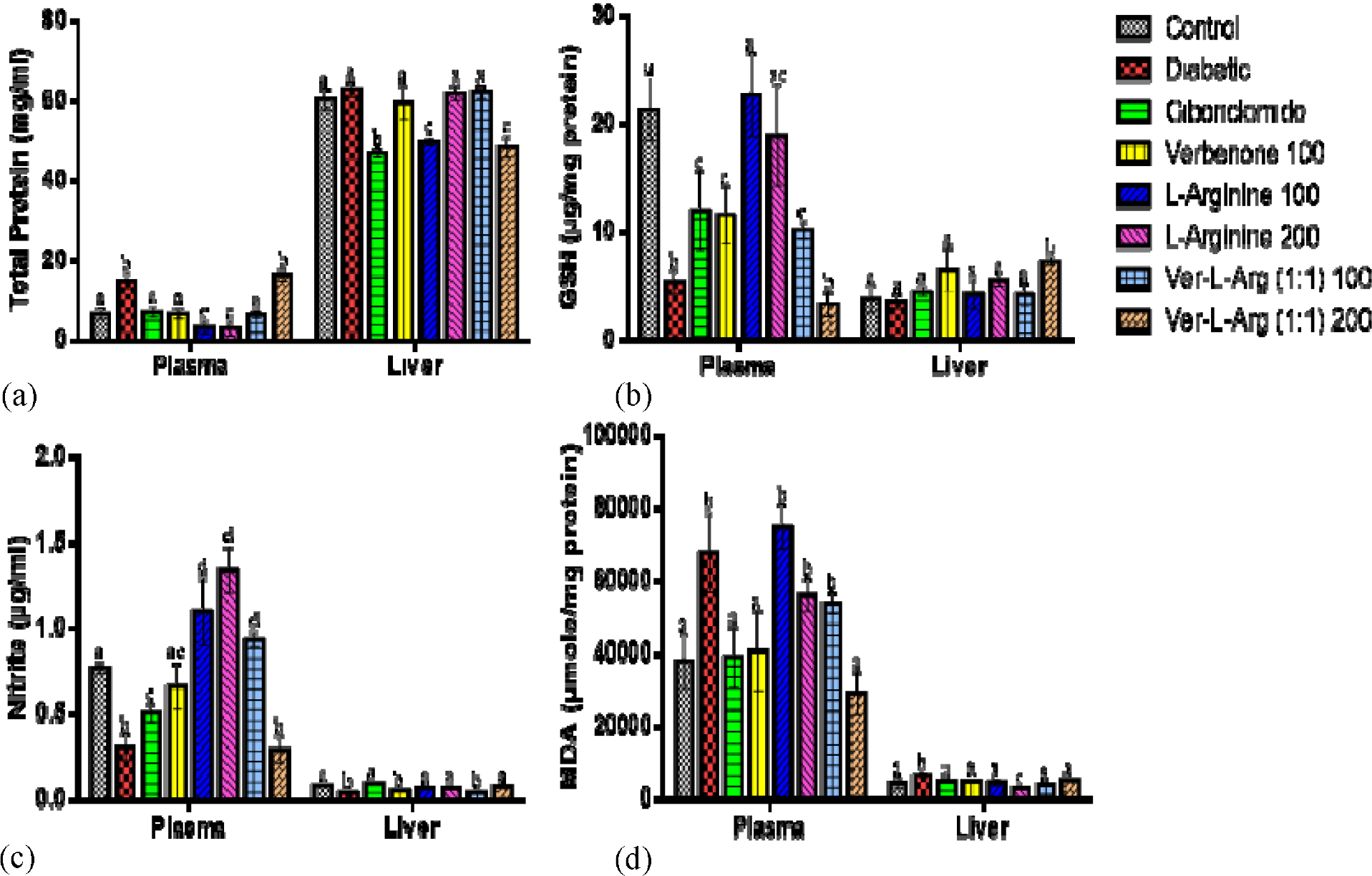
Total protein level and *in vivo* none enzymatic antioxidant markers in plasma and liver of diabetic mice treated with verbenone, L-arginine and their ratio combination. (a) Total protein level (b) GSH level (c) NO concentration (d) MDA level. Values are expressed as mean ± SEM, n = 5. Values with different superscripts are significantly different. Ver-L-Arg (1:1) = Verbenone + L-arginine (1:1 v/m)

Plasma nitrite (pNO) concentration was significantly decreased (p<0.05) in diabetic untreated, glibenclamide and Ver-L-Arg 200 groups compared with control group whereas, the level of pNO was significantly increased (p<0.05) in L-arginine 100, L-arginine 200 and Ver-L-Arg 100 groups (Figure 3c). In the liver, nitrite (lNO) concentration was significantly decreased (p<0.05) in diabetic, verbenone 100 and Ver-L-Arg 100 groups. Furthermore, significant increase (p<0.05) were observed in plasma and liver malondialdehyde in diabetic untreated, L-arginine 100, L-arginine 200 and Ver-L-Arg 100 groups compared with control group (Figure 3d). But, following treatment with the compounds, a significant reduction (p<0.05) was observed in glibenclamide, verbenone 100, and Ver-L-Arg 200 groups compared with control group. Similar effects were expressed in the liver, with a significant decrease (p<0.05) after treatment except in L-arginine at 200 mg/kg body weight.

### 3.4. Effects of verbenone, L-arginine and their combination (Ver-L-Arg) on enzymatic antioxidants indices

The activity of plasma GST (pGST) was significantly increase (p<0.05) in diabetic untreated, L-arginine 100, L-arginine 200, and Ver-L-Arg 100 groups compared with control group (Figure 4a). However, the glibenclamide, verbenone 100, and Ver-L-Arg 200 groups showed significant decease in the level of pGST compared to the diabetic untreated group. In the liver, significant increase (p<0.05) in the activity of GST was observed in all the treated groups after 14 days except in Ver-L-Arg 100 and Ver-L-Arg 200 groups. The activity plasma SOD (pSOD) was significantly decreased (p<0.05) in diabetic untreated, L-arginine 200, and Ver-L-Arg 100 groups. All the diabetic treated groups expressed significant increase (p<0.05) in the activity of liver SOD (lSOD) when compared with diabetic untreated group (Figure 4b).

**Figure 4.**
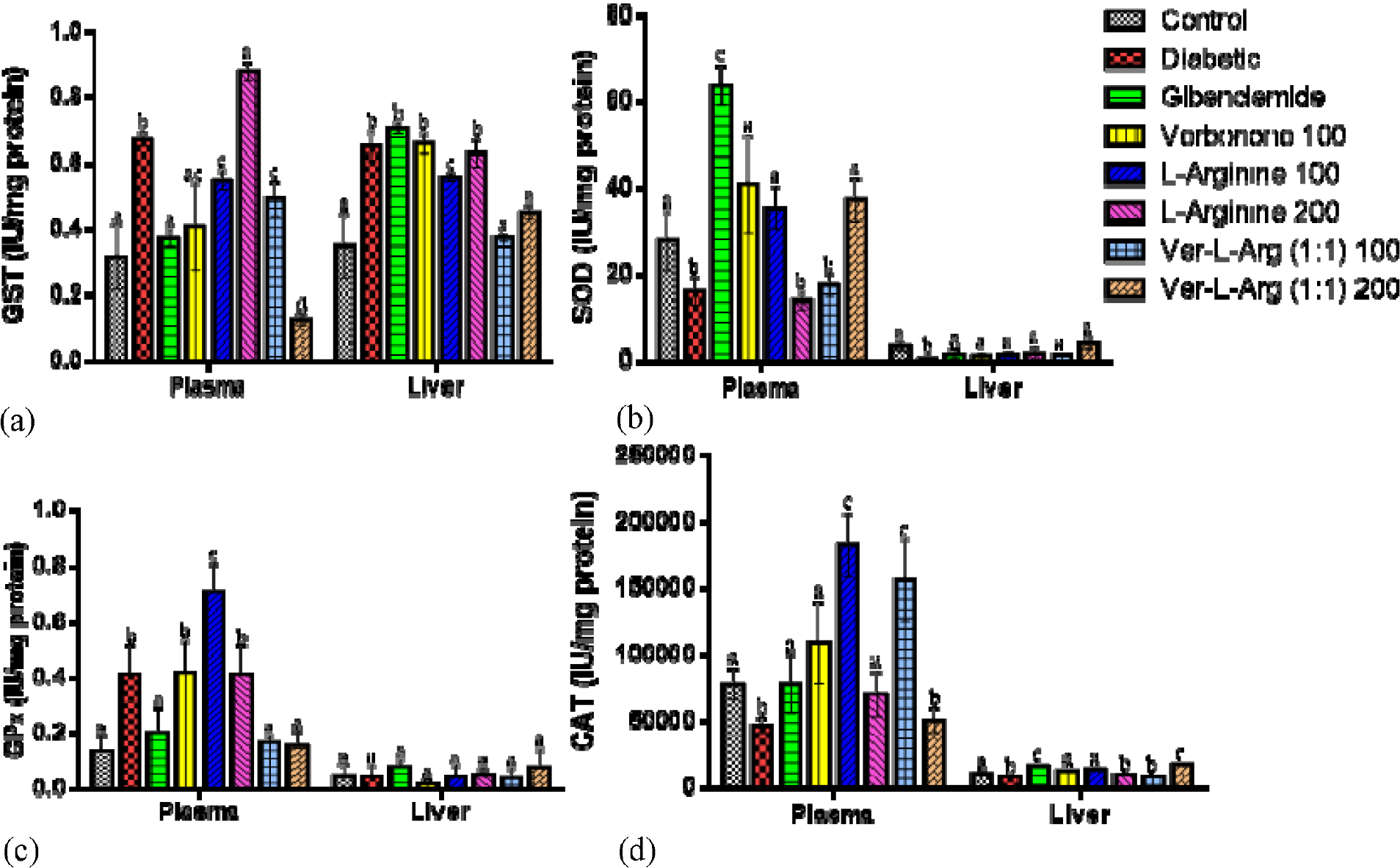
*In vivo* enzymatic antioxidant markers in plasma and liver of diabetic mice treated with verbenone, L-arginine and their ratio combination. (a) GST activity (b) SOD activity (c) GPx activity (d) CAT activity. Values were expressed as mean ± SEM, n = 5. Values with different superscripts are significantly different at p<0.05. Ver-L-Arg (1:1) = Verbenone + L-arginine (1:1 v/m)

Plasma GPx (pGPx) activity was significantly increased (p<0.05) in diabetic untreated, verbenone 100, and L-arginine 100 and L-arginine 200 groups. But, the activity of this antioxidant enzyme deceases in the glibenclamide, and Ver-L-Arg 100 and Ver-L-Arg 200 groups (Figure 4c). In the liver, GPx (lGPx) activity was not significant (p>0.05) in all treated groups compared with the control group. the activity of plasma CAT (pCAT) was significantly increased (p<0.05) in the glibenclamide, verbenone 100, L-arginine 100, L-arginine 200, and Ver-L-Arg 100 groups compared with diabetic untreated group. However, significant increase (p<0.05) in CAT activity was also observed in the liver after 14 days treatments, except in L-arginine 200 and Ver-L-Arg 100 groups compared with diabetic untreated group (Figure 4d).

### 3.5. Moelcular docking interactions of verbenone and L-arginine in antioxidant proteins

Verbeneone and L-arginine interacted with antioxidant proteins (Table 2). Verbenone interacted with lower binding energy (-6.0 Kcal.mol^-1^) compared with L-arginine (-4.3 Kcal.mol^-1^) and 5- fluorouracil (-4.9 Kcal.mol^-1^) in CP450. However, in LO, MP and NAD(P)HO the reference compound interacted with lower energies (-7.0, -6.6 and -8.0 Kcal.mol^-1^) compared with verbenone (-6.0, -5.3, -6.2 Kcal.mol^-1^) and L-arginine (-5.5, -5.3, -4.7 Kcal.mol^-1^) respectively. Furthermore, verbenone was superior in binding energy interactions compared with L-arginine. 5-fluorouracil interacted with ASP^349A^ (3.73Å), LYS^420B^(4.69Å), SER^343A^(3.24Å), ARG^342A^(5.02Å), ASP^414A^ (2.72Å); verbenone interacted with PHE^100B^(4.28Å), LEU^388B^(5.46Å), PRO^367B^(4.17Å), PHE^476B^(3.51Å) while L-arginine interacted with GLU^415A^(2.71Å), GLU^81B^(2.03Å), TYR^424B^(2.72Å), ASP^414A^(2.36Å) and LYS^423B^(3.66Å) in the binding pockets of human cytochrome P450 (CP450) (Figure 5a). Verbenone only interacted with PHY^155A^(4.81Å) while L-arginine interacted with TRP^791A^(3.59Å), LYS^545A^(2.58Å), PHE^162A^(2.32Å), VAL^144A^(2.14Å) in LO protein (Figure 5b). In MP, melatonin and L-arginine interacted with similar amino acids in the binding sites including ARG^333C^, ARG^239C^, and ASP^94A^ (Figure 5c). While dextromethorphan expressed lower binding energy compared with verbenone, and L-arginine, the reference compound also interacted with more amino acid residues (Figure 5d).

**Table 2.** Binding energies (Kcal.mol^-1^) of verbenone and L-arginine with some antioxidant proteins.

| Compounds | Binding energy (Kcal.mol<sup>-1</sup>) |  |  |  |
| --- | --- | --- | --- | --- |
|  | CP450 | LO | MP | NAD(P)HO |
| 5-fluorouracil (FLU, R1) | -4.9 |  |  |  |
| Zileuton (ZIL, R2) |  | -7.0 |  |  |
| Melatonin (MEL, R3) |  |  | -6.6 |  |
| Dextromethorphan (DEX, R4) |  |  |  | -8.0 |
| Verbenone | -6.0 | -6.0 | -5.3 | -6.2 |
| L-arginine | -4.3 | -5.5 | -5.3 | -4.7 |

Human cytochrome P450 (CP450), lipoxygenase (LO), human myeloperoxidase (MP), NAD(P)H oxidase (NAD(P)HO), R1 – R4 (References)

**Figure 5.**
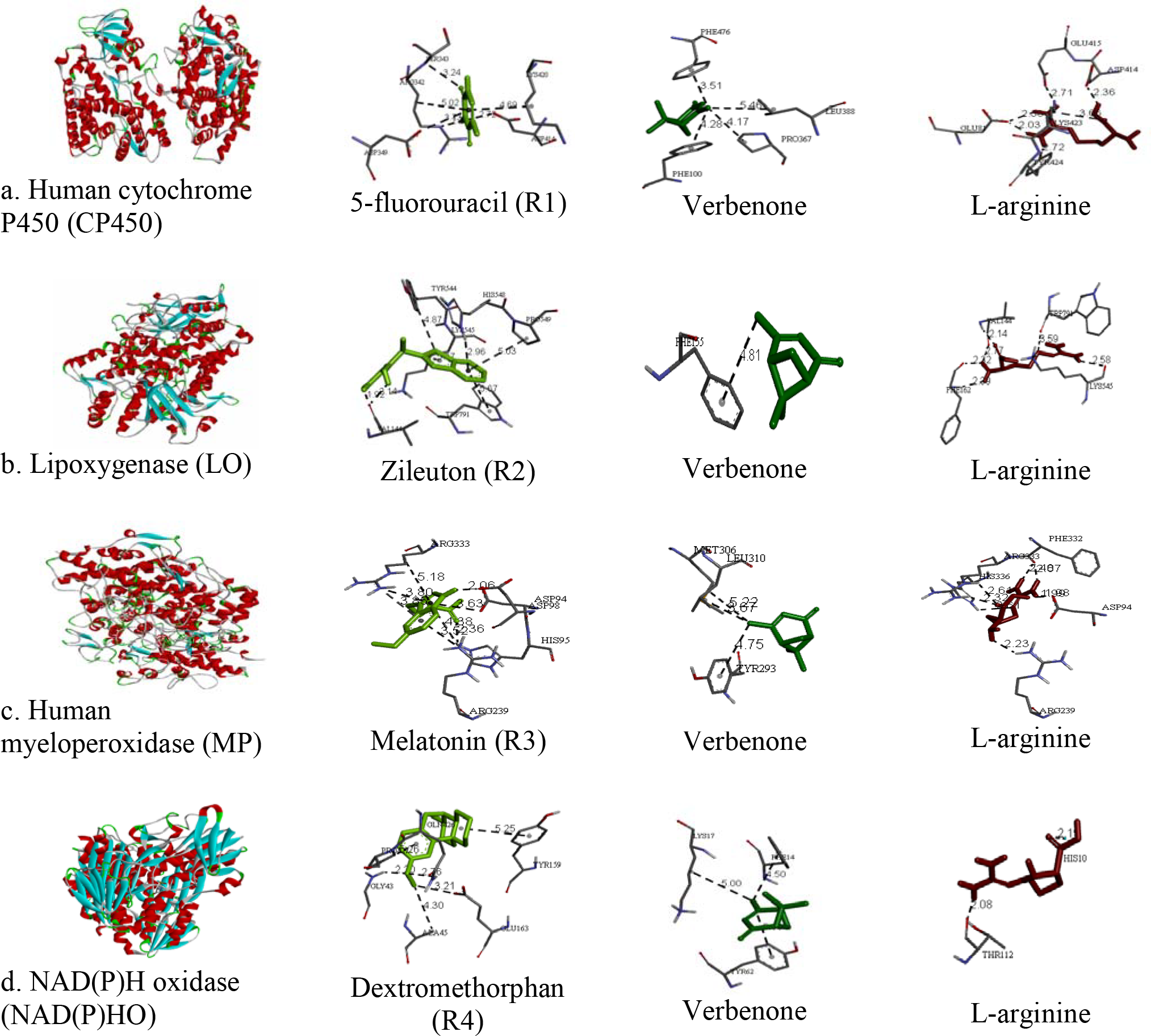
Docking poses of verbenone and L-arginine in binding pocket of antioxidant proteins. a. human cytochrome P450 (CP450), b. lipoxygenase (LO), c. human myeloperoxidase (MP), and d. NAD(P)H oxidase (NO), R1 – R4 (References).

## 4. Discussion

Free radicals are substances or atoms with one or more unpaired electrons. Their propagation brings about many adverse reactions leading to extensive tissue damage and the progression of diseases (Johansen *et al*., 2005). Lipids and proteins components of cells and tissues are susceptible to the attack of these free reactive radicals resulting to oxidative stress. However, antioxidants are compounds that prevent the damages on cells and tissues caused by reactive free radicals. The antioxidant activities of verbenone, L-arginine, and their ratios were demonstrated by their DPPH, NO, and H_2_O_2_ scavenging actions and TAC, and FRAP potentials. Verbenone demonstrated stronger NO and H_2_O_2_ scavenging properties compared with vitamin C. Even though, H_2_O_2_ is a weak oxidizing agent, it can inactivate a few enzymes directly, through the oxidation of essential thiol (-SH) groups (Thams and Capito, 1999). Moreso, it crosses cell membranes rapidly, reacting with Fe^2+^, and /or Cu^2+^ ions forming hydroxyl radicals leading to their toxic effects in cells (Dontha, 2016). Verbenone and L-arginine could acts by donating an electron to H_2_O_2_, then converting it to water and preventing their accumulation cells. The antioxidant activities were also demonstrated by the ratio combination of the verbenone and L-arginine. Thus, suggesting the potent antioxidant activities of the compounds.

Oxidative stress caused by reactive free radicals plays vital role in the development and progression of several diseases, including DM. But, the severity of oxidative stress-associated diseases can be attenuated by scavenging the reactive free radicals, therefore preventing their development and progression. Functional foods are soruces of natural antioxidants that can inhibits the detrimental actions of these reactive molecules. Therefore, diseases such as DM, which are associated with oxidative stress induced by radicals could be manage by antioxidants. In light of this, the management of DM is multifacet involving several management options (Tijjani *et al*., 2017). In the present study, plasma glucose level was significantly increased in the diabetic untreated mice following the IP injection of STZ, which is an indication of successful diabetes induction (Olatunde *et al*., 2021). Treatements with verbenone, L-arginine and their combination resulted in significant decrease in blood glucose level. Previous studies have documented the antidiabetic action of *D. carota* and some of its biocompounds (Khaki, 2011; Pouraboli and Ranjbar, 2015; Tijjani *et al*., 2022). The presence of the two compounds in *D. carota* seed could be partly responsible for the antidiabetic properties of the seed. The antioxidant activity of *D. carota* seed could also provide synergetic advantage of the seed in treatment of DM (Tijjani *et al*., 2019; Tijjani *et al*., 2020a; 2020b; 2020c; Tijjani and Imam, 2021) due to the antioxidant action of the verbenone and L-arginine. L-arginine admistration alone did not produce the desired blood glucose lowering effect compared to verbenone or glibenclamide. Thus, this suggests a synergetic effect of the co-administration of verbenone and L-arginine. Furthermore, the blood-reducing action of the two compounds could be linked to their antioxidant actions.

Peroxidation is an auto-catalytic and destructive process whereby polyunsaturated fatty acids in cell membranes undergo degradation to form lipid hydroperoxides (Linden *et al*., 2008). This reaction is caused by reactive oxygen species resulting to the formation MDA (Baliga *et al*., 2018). Subsequently, the reactive MDA causes toxic stress in cells. The elevation in the levels of pMDA and lMDA recorded in diabetic untreated mice may be due to the induction of oxidative stress in plasma, and thus lipid peroxidations, leading to increase pMDA production which was attenuated by verbenone and L-arginine. A similar effect displayed by *D. carota* seed, indicated that extract of the plant seed significantly reduced the concentration of MDA in hyperlipidemic mice, and was linked to the mop-up ability of formed radicals that can cause peroxidation, or prevent the accumulation of the peroxidation product, which could cause several harmful effects (Tijjani *et al*., 2020b).

The non-enzymatic antioxidant, glutathione is one of the most abundant tripeptides present in the liver. Its functions are mainly concerned with the removal of free radical species such as hydrogen peroxide, superoxide radicals, alkoxy radicals, and maintenance of membrane protein thiols and as a substrate for GPx and GST (Prakash *et al*., 2001). The significant increase in pGSH observed following the treatment of diabetic mice with verbenone and L-arginine treatments may be due to the potential of the compounds to increase cellular level of the non-enzymatic antioxidant. However, the non significant effects observed in lGSH except within the ver-L-arg 200 group may suggest that the removal of free radical species may not be adversely affected by the treatments rather it will enhance the antioxidant defense system in the experimental animals.

CAT, SOD, and GPx are important endogenous antioxidant marker enzymes for cellular enzymatic system (Zuo *et al*., 2012). SOD dismutates superoxide radicals forming hydrogen peroxide while CAT decomposes them and protects tissues from highly reactive hydroxyl radicals (Yao *et al*., 2005; Nandy *et al*., 2012). Verbenone played a vital role in the modulation of pCAT, by increasing the concentration compared with diabetic untreated mice. The two compounds elevated the activities of these antioxidant enzymes, thus indicating their potential in enhancing the scavenging properties, the enzymatic antioxidant and subsequent prevention of oxidative stress in the diabetic mice.

One of the characteristics features of DM is the inability of cells to utilize glucose for energy production. The body resort to the utilization of tissue protein as energy source leading to decrease in protein storage and reduced body weight (Montilla *et al*., 2004). The significant high level of total protein in liver could indicate dehydration or oxidative stress that causes protein to accumulate abnormally in DM. The reversal of this diabetic condition by verbenone, L-arginine and their ratio combination is an indication of their likely potentials in the prevention or amelioration of diabetes induced alterations such as total protein in diabetes

*In silico* study was used to assess the antioxidant activity of verbenone and L-arginine. The method used are important to study receptor-ligand interactions in the inhibition of antioxidant related enzymes (Gupta *et al*., 2018). The selected proteins included human cytochrome P450 (CP450), lipoxygenase (LO), human myeloperoxidase (MP), and NAD(P)H oxidase (NAD(P)HO) which are implicated in reactive oxygen species (ROS) produced during the metabolism (Dharmaraja, 2017; Costa *et al*., 2018). The reference compounds; 5-fluorouracil (FLU, R1), zileuton (ZIL, R2), melatonin (MEL, R3), and dextromethorphan (DEX, R4) used in the evaluation of *in silico* antioxidant potentials are compound commercially available in drug inhibitory activity against their respective receptors (Allegra et al., 2001; Gunes et al., 2006; Liu et al., 2009; Saul et al., 2017).

In human myeloperoxidase protein, melatonin and L-arginine interacted with similar amino acids in the binding sites including ARG^333C^, ARG^239C^, and ASP^94A^, expressing L-arginine potential antioxidant ability. Verbenone and L-arginine demonstrated favorable binding energies and amino acid interactions in the selected proteins compared with the reference compounds. Verbenone was superior in its interaction with human cytochrome P450 (CP450) compared with 5-fluorouracil and L-arginine. Verbenone similarly demonstrated superior interactions with all other proteins compared with L-arginine. Suggesting that the compound could inhibit ROS production cycle, reduce oxidative stress, and maintenance of redox balance (Dharmaraja, 2017).

## 5. Conclusion

Verbenone, L-arginine, and their ratio combinations possess *in vitro* antioxidant properties as well as reduced blood glucose levels in STZ-NAD-induced diabetic mice. The observed *in vitro* antioxidant activities of verbenone, L-arginine, and their ratio combination were complemented *in vivo*. Furthermore, the compound interacted *in silico* with antioxidant proteins, expressing their additional antioxidant potential.

## Abbreviations

DM: diabetes mellitus
DPPH: 1,1-diphenyl-2-picrylhydrazyl
FRAP: ferric reducing antioxidant power
H_2_O_2_: hydrogen peroxide
LO: lipoxygenase
MP: human myeloperoxidase
NAD(P)HO: NAD(P)H oxidase
NO: nitric oxide
STZ: streptozotocin
TAC: total antioxidant capacity
GSH: reduced glutathione
MDA: malondialdehyde
GST: glutathione-S-transferase
SOD: superoxide dismutase
GPx: glutathione peroxidase
CAT: catalase.

## Notes

### Competing Interest Statement

The authors have declared no competing interest.

